# Cell Type Specific Inference of Perturbations in Synaptic Communication with MultiNeuronChat

**DOI:** 10.1101/2025.11.11.687822

**Authors:** Gianluca Volkmer, Jens Hjerling-Leffler, Lisa Bast

## Abstract

Inference of differential neuron-neuron and neuron-glia communication from single nucleus RNA sequencing data is a powerful technique to uncover altered communication pathways when comparing groups, such as disease and control, developmental stages, or age groups, or different treatments. Yet, the communication changes are typically not identified on the cell type pair level, limiting its resolution in terms of answering biological and medical questions. Here we present MultiNeuronChat, a highly resolved framework that utilizes gene expression measurements of single cells from case-control single-cell/nucleus RNA-seq data sets together with an existing comprehensive database comprising cell adhesion molecules, gap junctions and synaptic transmission for the differential analysis of neuron-neuron and neuron-glia communication. In an in silico study, we show our method accurately and efficiently predicts known cell type-specific perturbations without summarizing communication scores across donor samples. Using a published single-nucleus RNA sequencing dataset, we highlight the sensitivity of our method by showcasing that MultiNeuronChat identifies both known and novel communication pathways in several cell type pairs in Alzheimer’s disease. Lastly, we highlight that MultiNeuronChat’s donor-specific communication score calculation can be utilized to inform patient stratification.

## Introduction

Fundamental brain functions that govern behavior, decision-making, perception, memories, emotions, speech, and language are generated by specific communication patterns between neuronal and glial cell types (Lovinger 2008; Haydon 2000). Many psychiatric and neurological disorders feature pathological cell-cell communication that symptomatically manifests as cognitive and behavioral impairments in patients. Knowing the exact cell-type and ligand-target pairs that are impaired in cell-cell communication would be useful in the identification of underlying disease mechanisms and could guide the development of targeted treatments for many brain disorders.

Amongst neurons, a highly specialized form of cell-cell communication is realized through synaptic transmission, i.e. electrical and chemical signals that are sent and received at synapses. The sending neuron releases ligands, such as neurotransmitters, from vesicles, which then bind to receptors on the receiving neuron to alter its function. In addition, both neurons and non-neuronal cells communicate through paracrine signaling, ion fluxes, neurotransmitters, cell adhesion molecules, and signalling molecules at neuronal synapses and gap junctions (Fields and Stevens-Graham 2002). Cell-cell communication can be controlled by regulating the gene expression of ligand-producing genes in sending cell types and target subunit genes in receiving cell types.

Advances in single-cell/nuclei omics now enable the sequencing of the transcriptome of cells from model tissues, nuclei from post-mortem human brain tissue of patients and neurotypical controls, cells of induced pluripotent stem cell-derived organoids, or cell cultures with cell-type resolution. Simultaneously, sequencing technologies are becoming increasingly affordable, allowing experiments to be scaled to larger cohorts, which results in comprehensive resources for transcriptomic alterations at the cell type resolution for an increasing number of brain disorders.

Recently, several computational methods have been developed for inferring cell-cell communication networks from scRNA-seq data (Dimitrov et al. 2022; Efremova et al. 2020; Zhao et al. 2023; Y. Zhang et al. 2021; Jin et al. 2021; Hou et al. 2020; Cabello-Aguilar et al. 2020; Jakobsson, Spjuth, and Lagerström 2021; Kumar et al. 2018; Cillo et al. 2020; Tsuyuzaki, Ishii, and Nikaido 2023; Raredon et al. 2022; Hou et al. 2020; Noël et al. 2021; Browaeys, Saelens, and Saeys 2020). As the only method to date specifically intended to analyze brain cell communication, NeuronChat (Zhao et al. 2023) elegantly models the neurotransmitter release through vesicles, and the target abundance, which are used to estimate a communication score for specific cell type- and ligand-target-pairs. While this method employs an agglomerative approach, it allows to identify which ligand-target pairs significantly differ between two groups. It however does not indicate which cell type pairs communicate significantly differently between the case and control groups for a given interaction pathway, and if this effect was driven by individual donor samples. Having such information would enable finding answers to many interesting and relevant biological questions.

Here, we present MultiNeuronChat, a differential cell-cell communication inference method designed explicitly for brain cells, which utilizes an expanded ligand-target database and models to compare donor- and cell-type-specific communication scores between two groups. We thoroughly tested MultiNeuronChat *in silico* and showcase its fast, robust and accurate performance. Applied to a case-control single-nucleus RNA sequencing dataset, MultiNeuronChat confirmed several known ligand-target pairs, linked them to sending and receiving cell types, and revealed patient heterogeneity with respect to cell-cell communication in Alzheimer’s disease.

## Results

### Highly resolved communication score calculation using an extended database

Group-level synaptic communication scores can be approximated from single-nucleus/cell RNA-sequencing (sn/scRNA-seq) data by taking into account the expression of genes coding for ligand proteins expressed in the sending cell type and genes coding for target proteins expressed in the receiving cell type, as previously shown (Dimitrov et al. 2022; Efremova et al. 2020; Zhao et al. 2023). The information on which genes’ expression is required to synthesize the subunit proteins of a specific ligand or target is stored in a database.

To build a comprehensive database containing neurotransmitters, receptors, and gap junctions, we created the MultiNeuronChat database (Methods, Fig. 1A, Table T1), which contains all entries from the NeuronChat database (Zhao et al. 2023) with additional substitutions of composite receptors from the CellPhone database (Garcia-Alonso et al. 2022; Efremova et al. 2020). Using this comprehensive database and the modeling approaches of Zhao et al. 2023, ligand and target abundances are then estimated from normalized and summarized sn/scRNA-seq data. To summarize each gene’s abundance, log-normalized counts are averaged with either mean, trimmed mean for [0.05,0.95] or [0.1,0.9] quantiles, or trimean (default) for each cell type, and donor. Subsequently, estimated ligand and target abundances are used to calculate highly resolved donor-specific communication scores per sending and receiving cell type, together with the ligand of the sending cell type, and the target of the receiving cell type (communication score tetrads per donor, Methods, Fig. 1B). These scores are then aggregated into two distinct empirical distributions, one per condition, where each donor-specific communication score is treated as an independent observation of the respective underlying distribution. To limit loss of power due to multiple comparisons, MultiNeuronChat prefilters the communication score tetrads according to the Wasserstein distance of their condition-specific score distributions across donors. To determine communication differences between two conditions, these shortlisted tetrads are then compared between the two conditions with a user-specified statistical distribution test, including multiple comparison corrections (Methods, Fig. 1C).

**Figure 1.**
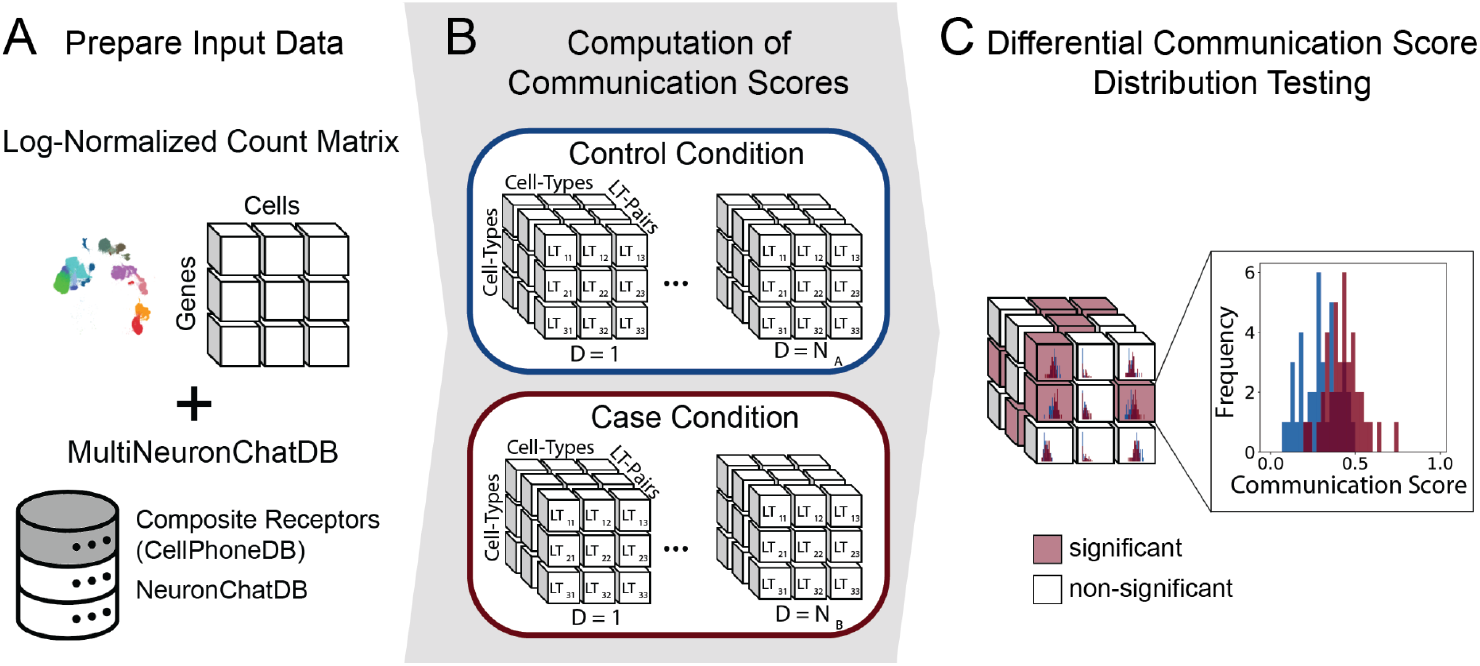
MultiNeuronChat methods overview. **A** First, Log-normalized counts from a case-control sc/snRNA-seq data set are averaged across cell types and donors for each relevant gene using optionally the mean, the trimmed mean for [0.05,0.95] or [0.1,0.9] quantiles, or the trimean. **B** Using the comprehensive MutliNeuronChat database (A) and averaged log-normalized counts, Donor-specific communication scores are then computed for each source cell type - target cell type - ligand - target tetrad. **C** Lastly, communication score distributions per condition are compared with a statistical test.

### Accurate and efficient identification of communication tetrads with largest differences between cases and controls

To assess the performance of MultiNeuronChat, we developed an *in silico* evaluation approach (Fig. 2A). By utilizing the snRNA-seq data simulation algorithm SPARSim (Baruzzo, Patuzzi, and Di Camillo 2020) and a snRNA-seq data set of human postmortem prefrontal cortex of neurotypical controls (Bast et al. 2025, Fig. S1A), we first estimated realistic parameters for the distribution of measurements for each sex and cell type cluster (Fig. S2A, Methods). According to these parameters and the estimated cell type composition (Fig. S2B), we generated an in silico case-control snRNA-seq dataset for each cell type cluster by applying predefined perturbations (Table T2) to the CASE group (Fig. 2A, S1B-D, S3, Methods). To further test scenarios in which only a subset of individuals in the CASE group obtain an increased or decreased expression for a specific ligand or target subunit coding gene, we varied the proportion of individuals in the CASE group that are affected by the perturbation (Fig. 2A,C).

**Figure 2.**
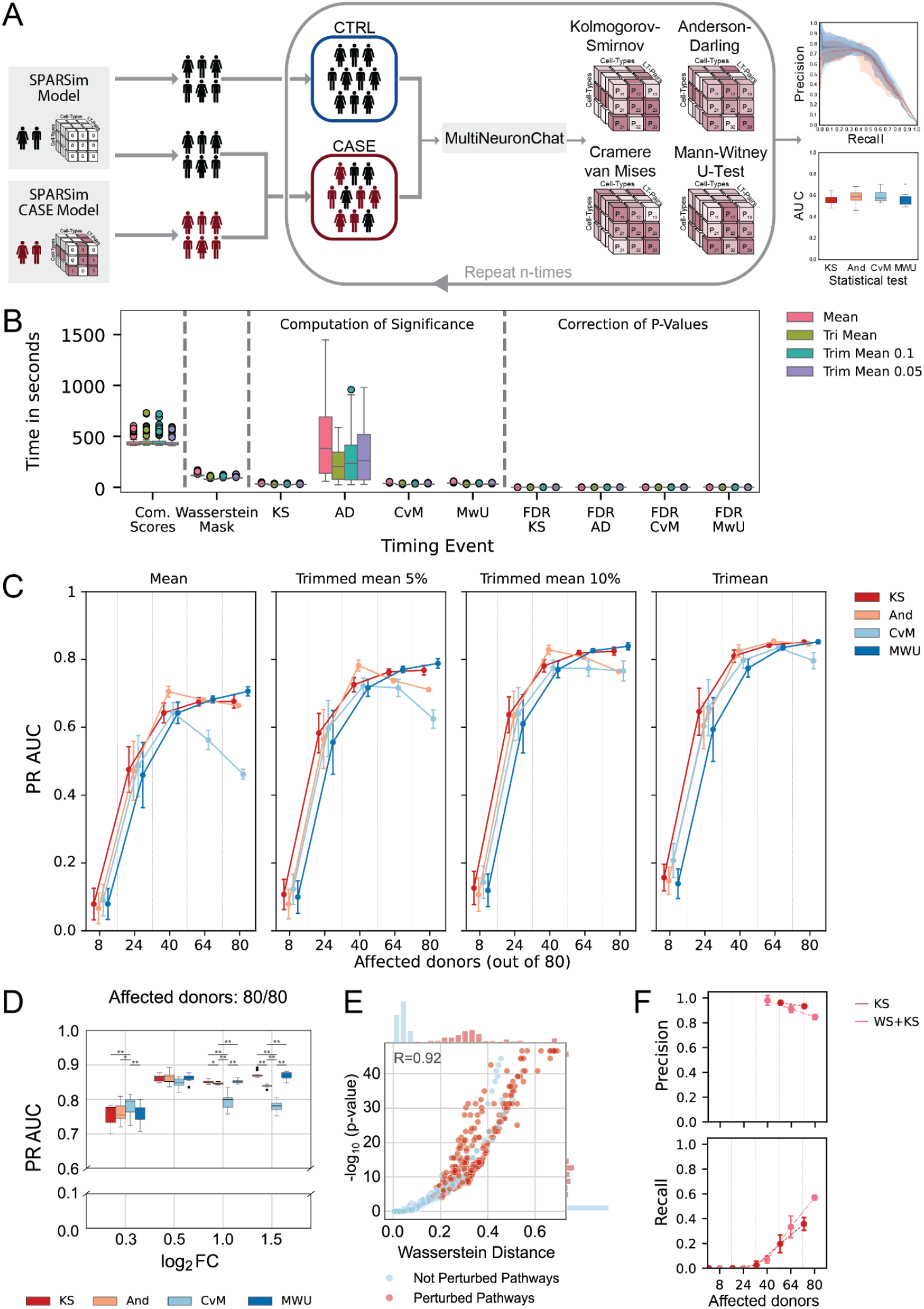
**A** In silico evaluation approach: SPARSim model with sex- and cell type-specific input parameters is utilized to simulate counts per cell type for male and female samples of an unperturbed group (SPARSim Model, black), and a perturbed group (SPARSim CASE Model, red). For the SPARSim CASE Model expression values of specific cell types and genes were systematically increased or decreased. CTRL group donors are randomly sampled from the unperturbed group. CASE group samples were randomly sampled from individual perturbed and unperturbed samples according to the fraction of affected donors. MultiNeuronChat uses four different averaging modes and four different statistical distribution tests. The performance is evaluated based on the AUC of precision-recall curves. **B** Runtime of MultiNeuronChat using 8 processes on our HPC server for all different mean-types and sub-tasks of the method pipeline. The mean runtime for the full MultiNeuronChat pipeline on our simulated dataset is between 9.53 and 14.42 minutes, depending on the statistical test used. **C** Comparison of the four different averaging modes evaluated based on their precision-recall AUC for a range of donors with the perturbation with a log2 fold-change of 1.0 for the four different statistical distribution tests. The trimean averaging performs overall best. **D** The Kolmogorov-Smirnov and Mann-Whitney U-test perform better than the other distribution tests when 100% of donors are affected by the perturbation and the effect size (log2 fold-changes in gene expression for the perturbations) ranges between 1.0 and 1.5. For the same scenario with a rather small effect size of 0.3, the Cramér–von Mises criterion outperforms the other distribution tests. **E** Wasserstein distance (p^w^=10) compared to negative log10 p-values of MultiNeuronChat with trimean averaging and Kolmogorov-Smirnov test for perturbations with an effect size of 1.0 affecting 100% of the donors. High Pearson correlation (R=0.92) suggests meaningful prefiltering of communication tetrads using Wasserstein distance (red: perturbed pathways, blue: non-perturbed pathways). **F** 8.68 percentage points precision decrease for each statistical distribution test when not filtering for the top 10% of Wasserstein distances as compared to exclusively applying the statistical test when perturbing 100% of case donors with an absolute log2 fold-change of 0.3 (top) and 21.28 percentage points improved recall when applying the Wasserstein distance prefiltering compared to exclusively applying the Kolmogorov-Smirnov test (bottom), for both p-values were corrected with Benjamini-Yekutieli.

Depending on the statistical test used, the overall runtime of MultiNeuronChat on our simulated dataset using 8 simultaneous processes on our HPC server was between 9.53 and 14.42 minutes (Fig. 2B, Table T3).

Evaluation of MultiNeuronChat on simulated data revealed that the trimean expression averaging performed better than the other averaging modes (Fig. 2C, S4, T4). As expected, the performance improved overall with an increasing number of affected donors in the CASE group and with increasing effect size of the introduced perturbation. When introducing a perturbation with a relatively high absolute log_2_ fold change (FC) of 1.0 or 1.5 affecting a majority of donors in the CASE group, the average AUC of the precision-recall curves for the trimean reaches values of 0.85. Even for small effect sizes (log_2_ FC = 0.3), the distribution tests detect the perturbations with relatively high PR AUC values of 0.76 if all donors are affected (Fig. S4A-B). Across different effect sizes, we observed a relatively good performance if the gene expression of the majority of donors in the CASE group was perturbed and only a minor further improvement if all 80 donors, instead of only 64 had the perturbation (Fig. S4A-C).

To compare the communication score distributions between the two conditions, we implemented and tested the performance of four different distribution tests: the Kolmogorov-Smirnov (KS) test, the Cramér-von Mises (CvM) criterion, the Anderson-Darling (And), and the Mann-Whitney U-test (MWU). The performance of these four tests varied less for the trimean or trimmed mean 10%, as opposed to the mean or trimmed mean 5% averaging modes, especially for larger proportions of donors affected (Fig. 2C). When using the best performing averaging mode (trimean, Fig. 2C, S3) and applying a range of effect sizes (log_2_ FC ∈ { 0.3, 0.5, 1.0, 1.5}) to perturbations affecting {8,24,40,64,80} donors, the Kolmogorov-Smirnov test performed best, followed by the Cramér-von Mises criterion, Mann-Whitney U-test, and Anderson-Darling test, according to the statistical test ranking (Methods, Table T4). Assuming all donors are affected, the Kolmogorov-Smirnov test performs better than the other distribution tests for larger effect sizes (log_2_ FCs in gene expression for the perturbations of and 1.5). For a rather small effect size of 0.3, the Cramér-von Mises criterion outperformed the other statistical distribution tests (Fig. 2D). As MultiNeuronChat performs the distribution test for each communication score tetrad, which results in a very high dimensional search space with many partly dependent hypotheses, the strict multiple comparison correction (Benjamini and Yekutieli 2001), which assumes arbitrary dependence, would restrict the applicability of the method. To extend the range of scenarios where our method can be applied, we sought to create a prior ranking of communication pathways to reduce the number of hypotheses tested. We investigated the Wasserstein distance, which is a similarity metric of two distributions (Kantorovich 1960; Ramdas, Garcia, and Cuturi 2015), and tested the performance of our method when using a statistical distribution test with and without a Wasserstein distance pre-filtering (Fig. 2E-F). For the best performing average mode (trimean), we observed a high correlation (Pearson’s R=0.92) of the Wasserstein distance (parameter *p*_*w*_ =10) with the log p-values of the best performing test (Kolmogorov-Smirnov test, Fig. 2E). Lastly, we quantified how much the pre-filtering of the top 10% communication score tetrads using the Wasserstein distance impacts the precision and recall for each statistical test. We found that the recall improved by 21 percentage points, while the precision only decreased marginally (9 percentage points) when applying the Wasserstein distance prefiltering (Fig. 2F) and an absolute log2 fold-change of 0.3. For an absolute log2 fold-change of 1.5, the additional filtering has a subtle effect on precision and recall, resulting in an overall improvement of MultiNeuronChat in terms of applicability and precision (Table T4).

### MultiNeuronChat predicts communication changes in Alzheimer’s disease

To explore the performance of MultiNeuronChat on real data, we applied our method to a snRNA-seq dataset comprising more than 1 Million nuclei from the dorso-lateral prefrontal cortex of 141 patients with Alzheimer’s disease and 150 non-cognitive impairment donors. MultiNeuronChat identified both neuronal and neuro-glia communication to be impaired in Alzheimer’s disease (Fig. 3A), as reported previously (Bandyopadhyay 2021). Filtering for the top 1% Wasserstein distances and subsequent statistical testing yielded the same interactions as testing without prior filtering. Most prominently were the on average reduced CCK-CCKBR (Cholecystokinin to Cholecystokinin B Receptor) communication from astrocytes, microglia, oligodendrocyte progenitors, oligodendrocytes, inhibitory and excitatory neurons to excitatory neurons in Alzheimer’s disease (Fig. 3B-C). CCK is a neuropeptide and satiety hormone that plays an important role in learning and memory (N. Zhang et al. 2024). Since low CCK levels in cerebrospinal fluid were found to be strongly associated with Alzheimer’s disease biomarkers such as CSF total tau and p-tau181, as well as Alzheimer’s disease-like changes in cognition, CKK has previously been proposed as a metabolic biomarker for Alzheimer’s disease (Plagman et al. 2019). Moreover, MultiNeuronChat identified the communication through the neuronal adhesion molecule coding genes NRXN1 and NLGN1/ NLGN4X involving excitatory cells and oligodendrocyte progenitors to be overall strengthened in Alzheimer’s disease. This also includes homotypic communication, i.e., communication between excitatory neurons and between oligodendrocyte progenitors. Similarly, our method identified pronounced communication changes through NRXN3 and NLGN1/NLGN4X from inhibitory cells to excitatory cells and oligodendrocyte progenitors in Alzheimer’s disease (Fig. 3B-C). Interestingly, with a minor allele frequency of less than 1%, NLGN4X is a rare risk gene for Alzheimer’s disease (Belloy et al. 2024). Changes in neurexin-neuroligin have been implicated in altering the excitatory-inhibitory neurotransmission balance, which can lead to synapse damage and loss - one of the hallmarks of Alzheimer’s disease (Sindi, Tannenberg, and Dodd 2014; Serrano-Pozo et al. 2011).

**Figure 3.**
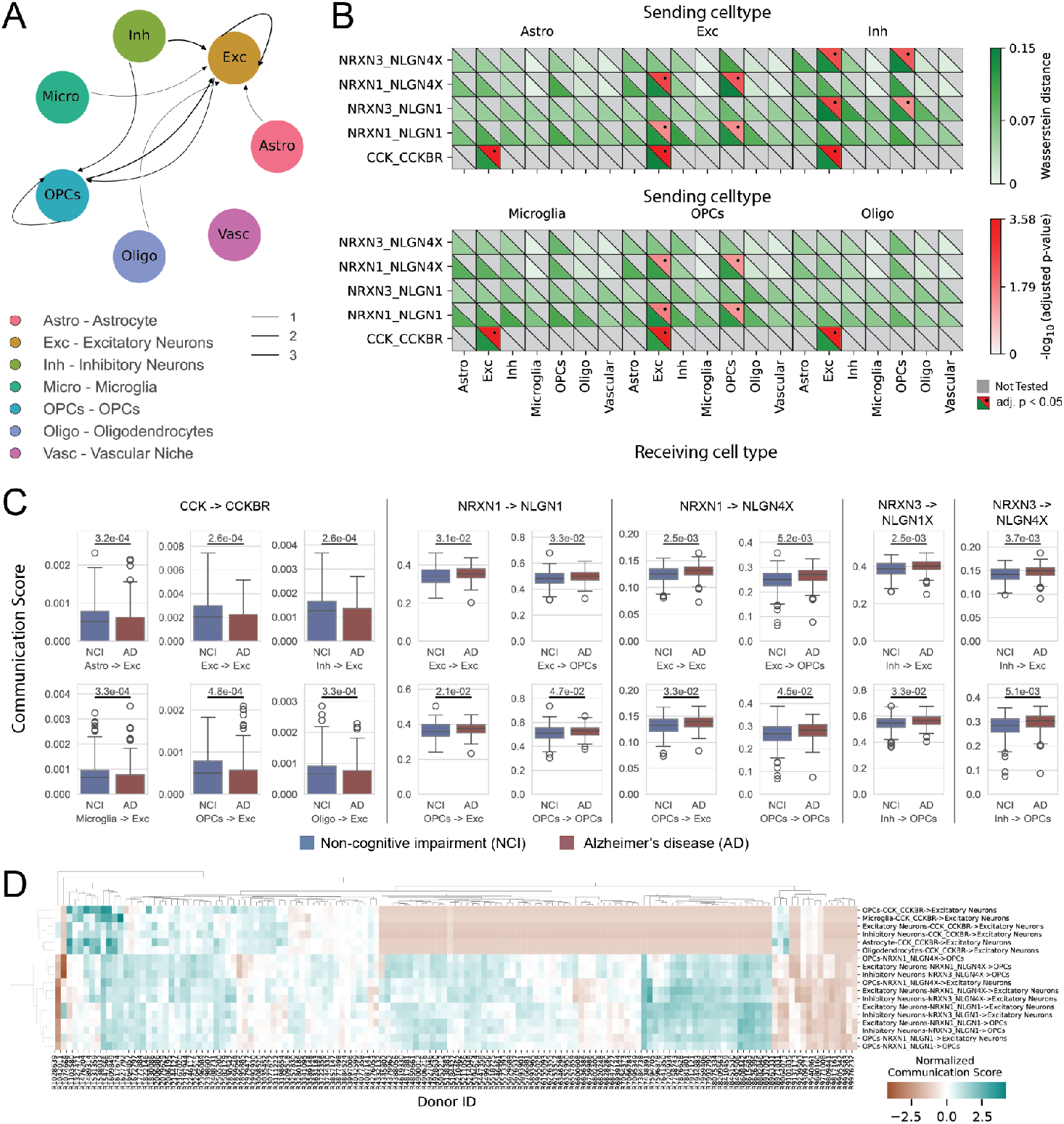
MultiNeuronChat applied to an Alzheimer’s disease-control snRNA-seq data set. **A** Cell type interactions with altered ligand-receptor interactions in the top 1% of Wasserstein distances/ with at least one significant interaction (alpha = 0.05, Kolmogorov-Smirnov test, p-values corrected with Benjamini-Yekutieli). **B** Aula-medica plot illustrating the negative log10 of adjusted p-values (red, upper triangles; dots mark significance) and Wasserstein distance (green, lower triangles) for ligand-receptor interactions with at least one cell-type pair in the top 1% of Wasserstein distances. Strong indications for perturbations in the CCK-CCKBR (Cholecystokinin to Cholecystokinin B Receptor) interactions between astrocytes/ microglia/ oligodendrocyte progenitors/ oligodendrocytes/ neurons and excitatory neurons, interactions involving NRXN1 and NLGN4X/ NLGN1 between excitatory cells and oligodendrocyte progenitors and interactions involving NRXN3 and NLGN4X/ NLGN1 between inhibitory cells and oligodendrocyte progenitors. **C** Communication score distributions per group for the 18 significant tetrads show overall significantly increased communication in Alzheimer’s disease, except decreased CCK-CCKBR communication (p-values of Kolmogorov Smirnov test). **D** Normalized communication score per Alzheimer’s disease donor for each significant tetrad to define communication patient subgroups. Communication scores were normalized to mean and standard deviation of NCI donor’s tetrad-specific communication scores (see Methods).

Lastly, we investigated MultiNeuronChat’s donor-specific communication scores to determine which communication changes co-occur across donors, and which ones occur in subgroups or individual donors. For the majority of Alzheimer’s disease patients, the CCK-CCKBR communication changes (upper block in Fig. 3D) and the NRXN1/3-NLGN1/4X communication changes (lower block in Fig. 3D) were each in the same direction, except for a few donors. We could identify five subgroups: donors with increased CCK-CCKBR and NRXN1/3-NLGN1/4X communication, donors with weak CCK-CCKBR and NRXN1/3-NLGN1/4X communication changes, donors with decreased CCK-CCKBR and increased NRXN1/3-NLGN1/4X communication, a small group of donors with increased CCK-CCKBR and decreased NRXN1/3-NLGN1/4X communication, and donors with decreased CCK-CCKBR and NRXN1/3-NLGN1/4X communication (Fig. 3D).

## Discussion

In this study, we evaluate and showcase MultiNeuronChat, the first comparative method for brain cell communication changes with donor- and cell-type-resolved communication scores. MutiNeuronChat’s database comprises cell adhesion molecules, gap junctions, and synaptic transmission for the differential analysis of neuron-neuron and neuron-glia communication. We have shown that our method efficiently computes communication scores, is accurate, and is based on robust statistics (AUC of precision-recall curves: 0.85). Compared to previously published methods, MultiNeuronChat models and computes highly resolved communication scores per cell type and donor. This allows for performing a statistical test on distributions across donors to identify cell type pairs with their ligand target pairs, identifying changes between a case and control group, rather than relying on mean values across cohorts, and thus circumvents error-prone averaging. Moreover, MutliNeuronChat outputs the exact cell types for which the identified cell-cell communication change occurs, and is at least 6-times faster than previous efforts. Our method enables a detailed investigation of brain cell communication alterations that can be applied to infer biological mechanisms between developmental stages, disease mechanisms, and inform patient stratification.

Using a large-scale snRNA-seq data set from Alzheimer’s patients and non-cognitive impairment control samples, we showcased MultiNeuronChat’s performance and usability on snRNA-seq data. MultiNeuronChat identified altered communication involving Cholecystokinin, a neuropeptide important for learning and memory, which has previously been associated with Alzheimer’s disease (Plagman et al. 2019, N. Zhang et al. 2024). Most importantly, MultiNeuronChat could pinpoint the CCK-CCKBR communication change to excitatory neurons as receiving cell types and reveal that approximately 60% of the donors have decreased CCK-CCKBR communication, 15% no CCK-CCKBR impairment, and 25% increased or mildly increased CCK-CCKBR communication. In addition, our method identified NRXN1/3-NLGN1/4X communication involving excitatory cells and oligodendrocyte progenitors in Alzheimer’s disease. Neuroligin has previously been implicated with memory impairment and Alzheimer’s disease (Arias-Aragón et al. 2023); (Dufort-Gervais et al. 2020; Tristán-Clavijo et al. 2015). The NRXN3-NLGN1, but not the CCK-CCKBR interaction, has been identified to be altered in Alzheimer’s disease by applying NeuronChat to an independent, much smaller (four case, four control samples) snRNA-seq data set from human postmortem entorhinal cortex tissue (Bartas et al. 2025).

A limitation of our approach is that MultiNeuronChat relies on comprehensive, high-quality input data, which however, is becoming increasingly available. It requires ligand and target coding genes to be expressed in many cells within the tissue of interest, and large sample sizes in both groups, especially for diseases with substantial patient heterogeneity regarding the expression of ligand and target coding genes. To accurately calculate the communication score approximations, MultiNeuronChat ideally requires a dataset with numerous reads per cell and a large number of cells per cell type per donor. To obtain the default setting of at least 10 cells per cell type per donor in the input data, a lower cell-type annotation resolution is at times necessary, compromising on the results’ resolution. Moreover, for the *in silico* evaluation, simulating snRNAseq data is not ideal since it does not fully capture the complexity of measurements. The database could be further improved by including the gene proportions required to build the subunits of ligands and targets. To inform this database extension, experiments in wildtype mice and healthy human cell or brain organoid cultures could be designed and utilized in the future. To inform the cell-cell communication even better, other omics data, such as transcriptomic reads measured from synaptosomes (Niu and Zong 2024; Chen et al. 2025), spatial transcriptomics, proteomics, and epigenomics could be incorporated, and the mathematical model possibly extended for this purpose. Upon availability, mutation status, and other patient-level data could be mapped against donor-specific scores to inform patient stratification tasks more comprehensively.

In summary, we present MultiNeuronChat, a computational method for the cell type specific inference of perturbations in synaptic communication from single-cell RNA sequencing data. We have shown that our method uses a comprehensive database, is accurate, efficient, and robust, and can reveal known and novel cell-cell communication differences between two groups. Since it provides the user with cell type and donor-specific scores, it is suited to investigate how many and which samples of the case group were affected by the inferred communication changes.

## Methods

### MultiNeuronChat

#### Database

The NeuronChat database (Zhao et al. 2023) forms the basis of our receptor-ligand interaction annotations, providing a well-curated resource for analyzing communication networks in both mouse and human contexts. However, NeuronChat (Zhao et al. 2023) does not include complex receptors, which are essential for accurately modeling many signaling pathways.

To address this, we extended the human database by incorporating data from CellPhoneDB (Garcia-Alonso et al. 2022; Efremova et al. 2020), a database containing several complex GABAA-, Kainate-, and NMDA-receptors. This integration has resulted in the “human_extended” database, combining the strengths of both resources to provide a more comprehensive and biologically relevant tool for cell-cell communication analysis in humans. Recognizing the dynamic nature of ligand-receptor interaction research, the database has been constructed to support future extensions. Researchers are encouraged to adapt the database to include novel interactions or emerging data as needed, ensuring its continued utility across various biological contexts.

#### Data normalization

Let *C* denote the raw count matrix of dimension *I* × *J*, where *I* is the total number of measured genes, and *J* is the total number of measured cells. The normalization process was adapted from (Zhao et al. 2023) to reflect our subject specific approach, and involves the following steps:

##### 1. Log-normalization of each cell

For each cell, the raw counts are normalized by dividing each entry *C*_*i,j*_ by the sum of all gene counts for the cell, scaling by 10,000, adding a pseudo-count of 1, and applying the natural logarithm

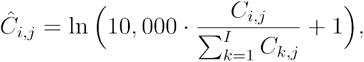

where *i* ∈ {1, …,*I* } is the gene index and *j* ∈ {1, …,*J* } is the cell/nuclei index.

##### 2. Subsampling genes

Following the log normalization, we exclude all genes that do not contribute to ligand or receptor synthesis, according to the MultiNeuronChat database. This will yield a new matrix Ĉ ^′^ ∈ *I*^′^ × *J*, where *I*^′^ ⊆ *I* is the subset of genes described in the database.

##### 3. Normalization across donors

The count matrix consists of *S* donors, where *S = N*_*A*_ + *N*_*B*_, with *N*_*A*_ donors from condition *A* and *N*_*B*_ donors from condition *B*. Let Ĉ ^′*s*^ be the log-normalized counts matrix containing only cells/ nuclei of donor *s* ∈ {1, …,*S* }. In a second normalization step, each donor-specific log-normalized counts matrix Ĉ ^′*s*^ is normalized by its maximum value:

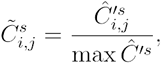

where max Ĉ ^′*s*^ denotes the maximum value of 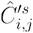 across *i* all *j* and for a specific donor *s*.

The two-step normalization ensures that technical variations are addressed within cells, while donor-level normalization ensures comparability among subjects.

### Inference of differential cell-type pair specific communication

Next, we compute the communication scores using the max normalized count matrix 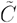.

#### 1. Computation of average gene expression

Let Ψ be the set of cell-type label annotations and let *K*^*s*^ ⊂ *J*^*s*^ be the subset of cells with indices belonging to cell-type *c* for donor *s*. For each donor *s* ∈ *S*, cell-type *c* ∈ Ψ, and gene *i* ∈ *I*^′^, we compute the average gene expression 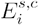 across cells/ nuclei using one of the following averages:

##### a Tri-mean expression

The tri-mean is a robust measure of central tendency that combines the median *Q*_2_, with the lower quartile *Q*_1_ and upper quartile *Q*_1_:

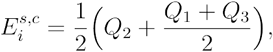

where *Q*_1_, *Q*_2_, and *Q*_3_ are computed from 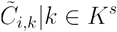.

##### b Arithmetic mean expression

The arithmetic mean provides a straightforward measure of central tendency:

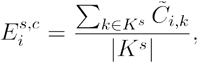

where |*K*^*s*^| is the cardinality (number of cells) of *K*^*s*^.

##### c Trimmed mean expression

The trimmed mean is similar to the arithmetic mean, but excludes the top and bottom *X* % of data points to remove the impact of the most extreme values, including outliers:

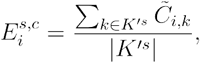

where *K*^′*s*^ ⊂ *K*^*s*^ excludes the top and bottom *X* % of values from *K*^*s*^.

#### 2. Computation of communication score matrices

On the molecular level, neuronal cell-cell communication is realized through the production and release of ligands/ neurotransmitters in the sending cell and the production of receptors/ targets in the receiving cell. To produce ligands and receptors, certain proteins need to be translated from the mRNA transcripts. To estimate the communication score of two given cell types from single cell/ nuclei RNA-sequencing data, the ligand- and receptor-abundances need to be computed.

Following the approach from (Zhao et al. 2023), the ligand- and receptor-abundances per *s* subject and cell-type *c* can be defined as following:

##### a Ligand abundance estimation

Let *m*_1_ be the number of distinct catalyzing steps of a particular ligand *i* and *P*_*m*_ the number of isoenzymes, i.e. enzymes that produce the same product, of step *m* ∈ [1, …,*m*_1_]. Let 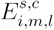 be the average expression of the *l*-th isoenzyme of catalyzing step *m* for ligand *i*, subject *s*, and cell-type *c*. If applicable for the specific ligand *i*, 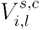 let be the average expression of the *l*-th isoenzyme that is needed for the synthesis of the vesicle that transports the ligand *i* across the membrane. The ligand-abundance *L*_*i*_ for a specific subject *s* and cell type *c* is then defined by

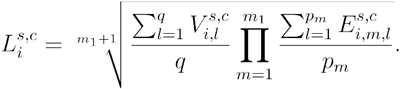

##### b Receptor abundance estimation

Let *m*_2_ be the number of distinct subunits the receptor *j* consists of and let denote *c*_*l*_ the stoichiometric coefficient for each subunit *l* ∈ [1, …,*m*_2_]. The abundance of the *j*-th receptor *R*_*j*_ is calculated as the geometric mean of the average expression of the subunits *E*_*j,m*_ raised to the power of their stoichiometric coefficients *c*_*l*_

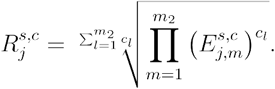

##### c Communication score tetrad estimation

For each known tetrad, i.e. ligand-receptor interaction (*i,j)* and cell-type pair (*c*_1_,*c*_*2*_) the communication scores are calculated by

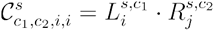

where the ligand abundance 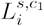 is taken from the source cell-type *c*_1_ and the target-abundance 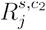 is taken from the receiver cell-type *c*_2_.

All interactions of a specific subject *s* are then represented as a communication score matrix 𝒞^*s*^ with dimensions | Ψ | × | Ψ | × | ℐ |, where is the number of cell types and | ℐ | is the number of known ligand-receptor interactions described in the MultiNeuronChat database. In this step, two sets of communication score matrices (one set per condition) 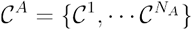and 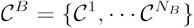 are calculated.

#### 3. Calculation of significantly different communication

Let 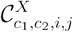 be the set of communication scores for the cell-type pair (*c*_1,_ *c*_2_) and ligand-receptor interaction for (*i,j*) all subjects of condition *X*, and 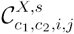 be the score for a specific subject *s* of condition *X*.

Interpreting 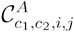 and 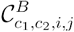 as collections of random variables from the underlying generating distributions 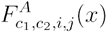 and 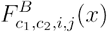, we can define the null and alternative hypothesis as:

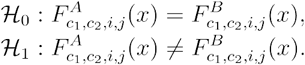

To test whether we should reject the null hypothesis one of the following four statistical tests is applied:

a. two-sample Kolmogorov-Smirnov test,
b. two-sample Anderson-Darling test,
c. two-sample Cramér–von Mises criterion,
d. two-sample Mann–Whitney U test.

In cases where the analytical implementation yields an undefined or zero p-value, or when a more precise estimate is explicitly requested, MultiNeuronChat falls back to a permutation-based computation of the p-value, using the respective test statistics, to obtain a more reliable estimate.

Finally, to correct for multiple hypothesis testing without assuming the independence of hypotheses, the Benjamini-Yekutieli procedure is applied.

##### Ranking of hypotheses using the Wasserstein metric

To address the challenge of testing all possible hypotheses, we introduce a ranking method based on the Wasserstein *p*-distance (Kantorovich 1960; Ramdas, Garcia, and Cuturi 2015). This approach prioritizes hypotheses by quantifying the difference between the empirical distributions of communication scores observed under two conditions. Specifically, we define the distribution *μ* to represent communication scores from condition *A* and *v* to represent scores from condition *B*.

The Wasserstein *p*-distance offers an intuitive metric for comparing distributions by assessing the minimal “cost” of transforming *μ* into *v*. Unlike standard metrics, such as mean or variance, the Wasserstein distance captures differences in the overall shape of the distributions, making it particularly sensitive to subtle shifts that may reflect biologically meaningful changes.

By ranking hypotheses using this distance, we allow reducing the number of tested hypotheses, limiting the necessary adjustment of multiple hypotheses testing, while retaining those most likely to exhibit significant differences. This pre-selection ensures that subsequent significance tests focus on hypotheses with the greatest potential biological relevance, improving efficiency and interpretability.

### Method evaluation

#### Strategy and evaluation metrics

To evaluate the performance of MultiNeuronChat, we developed an in silico evaluation pipeline. Using the dataset from (Bast et al. 2025), we first fitted two distinct models for each cell type: (1) a cell-count model, and (2) a transcription model based on SPARSim (Baruzzo, Patuzzi, and Di Camillo 2020). With these models, we simulated data for 60 male and 60 female donors for each of the following three groups: (i) a control group with no gene expression perturbations; (ii) a baseline perturbation group without actual perturbations, serving as a control for dilution effects; and (iii) several case groups featuring perturbations in the expression of genes involved in cell-cell signaling using several different log2 fold-changes. Perturbed genes and cell types were selected through a constrained random sampling strategy, ensuring only relevant, actively expressed genes were included.

To assess method sensitivity, datasets were generated across varying magnitudes of log2 fold-changes (0.3, 0.5, 1.0, and 1.5) and proportions of affected donors (0.1, 0.3, 0.5, 0.8, and 1.0). For each combination of these parameters, we produced 10 replicate datasets, by randomly sampling from the previously simulated donors, to quantify variability in performance. After processing these datasets with the MultiNeuronChat pipeline, we evaluated the results using Precision-Recall curves and calculated the Area-Under-the-Curve (AUC). These metrics effectively handle imbalanced class distributions, which is essential for our scenario, as only a small subset of cell-cell interactions were perturbed. Additionally, we computed precise values for precision and recall following hypothesis filtering based on the Wasserstein metric, defining true perturbations as those with computed p-values below 0.05.

### Fitting a single-nuclei RNA-sequencing model

To evaluate our method *in silico* we required a simulated snRNA-seq dataset with and without perturbations. To achieve this, we first fit two models on the control group of Bast *et al*. 2025 paper:

1. a cell-count per cell-type model and
2. a gene expression model.

To fit a cell-count model, we first split the dataset into their respective cell types. For each cell-type *c* we then fit a gamma distribution model 𝒯_*c*_ (φ_*c*_,*Y*_*c*_) on the number of observed cells per donor, where φ_*c*_ is the shape and *Y*_*c*_ is the scale parameter of the gamma model.

For the gene expression model we utilize SPARSim (Baruzzo, Patuzzi, and Di Camillo 2020), a Gamma-Multivariate Hypergeometric model which has been shown to perform well in fitting general data properties while also preserving biological signals (Cao et al. 2021). As there exist certain genes that are sex-specifically expressed, we first split the control group into male and female subclasses. Following this, we fit cell-type specific SPARSim (Baruzzo, Patuzzi, and Di Camillo 2020) models. Each sex- and cell-type-specific model 𝒮_*s,c*_ (G,L) consists of a set of tuples G = { (Φ_*i*_,*Z*_*i*_) | *i* ∈ {1, …, *I* } }, describing the shape Φ_*i*_ and scale *Z*_*i*_ gamma distribution parameters for each gene *i* ∈ { 1, …,*I* } and a set of observed library sizes L = {*l*_1,_ …,*l*_*J*_}, where *J* is the number of cells with cell-type *c*, which are used for the Multivariate-Hypergeometric distribution.

#### Sampling of communication pathways to perturb

To evaluate our developed method, we randomly sampled a set of 306 pathways that we define as perturbed. A pathway is a tetrad (*s,r,l,t*) describing the affected source cell-type *s*, receiver cell-type *r*, ligand *l* that is produced in *s*, and receptor *t* that is produced in *r*.

We define two sets of possible perturbations:

1. source-ligand pairs (*s,l*) and
2. receiver-receptor pairs (*r,t*).

Each pair in either category was eligible to be randomly drawn as a perturbation, if the SPARSim (Baruzzo, Patuzzi, and Di Camillo 2020) model, that was fitted on the control group of (Bast et al. 2025), predicted non-zero expression for all production-steps of the corresponding ligand *l* in a cell-type *s* or non-zero expression of all subunits of a specific receptor *t* in cell-type *r*.

When a source-ligand pair (*s,l*) is drawn from the set of possible perturbations, all associated pathways (*s,r*_*i*_,*l,t*_*j*_) are defined to be changed, where *t*_*j*_ are all receptors that interact with ligand *l* and *r*_*i*_ are all receiver cell types that show expression of *t*_*j*_. Similarly, when a receiver-receptor pair (*r,t*) is drawn, all pathways (*s*_*i*_,*r,l*_*j*_,*t*)are defined to be perturbed.

We sampled from these pools of available source–ligand (*s,l*) and receptor–receiver (*r,t*) combinations until a minimum of 300 distinct pathways were affected, removing any ligand or receptors and its associated counterparts from the pool once it was selected, which ensures non-overlapping or redundant choices.

Once we have sampled a list of all source-ligand pairs (*s,l*) and receiver-receptor pairs (*r,t*) to perturb, we then sample a single gene to be either up or down regulated from the ligand production chain or set of receptor subunits.

The final list of perturbed pathways can be found in Table T2.

#### Simulating a new sample

To simulate a new sample *d* with sex *s*, we employ the following procedure:

For each cell-type *c* ∈ {1,…,*C* } we first sample the number of cells to simulate:

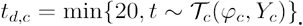

Then we sample a set of library sizes with the size *t*_*c*_ from the in the fitting procedure observed library sizes L_*c*_:

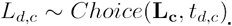

If we want to simulate a perturbed expression of a specific gene *i* by a log2 fold-change *p*, we adjust the corresponding scale parameter *Z*_*i*_:

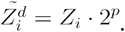

Using the original or adjusted parameters, we then simulate a gene expression matrix 𝒞_*c*_ of dimensionality *I* × *t*_*c*_ using SPARSim (Baruzzo, Patuzzi, and Di Camillo 2020):

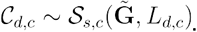

Once all cell-type specific expression matrices have been simulated for the individual *d*, they are combined into a single expression matrix:

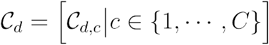

To simplify later notation, let us denote this whole simulation procedure for a single individual *d* with sex *s* as

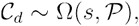

where 𝒫 = { (*c*_1_,*i*_1_,*p*_1_), …, (*c*_*n*_,*i*_*n*_,*p*_*n*_) } denotes the set of triplets defining the log2 fold-changes *p*_*x*_ that should be applied to gene *i*_*x*_ in cell-type *c*_*x*_.

### Creating the in silico dataset

We simulated several sets of samples, each consisting of *n* = 60 donors of sex *s* with different log2 fold-changes λ_0_ = λ_1_ = 0, λ_*i*∈{2,3,4,5}_ ∈ {0.3, 0.5, 1, 1.5}

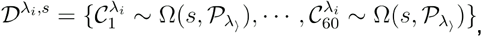

With 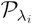 being the set of gene perturbations with log2 fold-change λ_*i*_. In the case of the control group or the baseline case group, with λ_0_ = λ_1_ = 0, the set of perturbations is the empty set 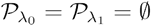.

In order to evaluate how sensitive our method is regarding both the effect-size of the gene expression change, and number of donors with the perturbation, we sample per log2 fold change λ_*i*∈{2,3,4,5}_ datasets 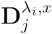 with *x* ∈ {10, 30, 50, 80, 100 }\% of case donors effected by the defined gene perturbations:

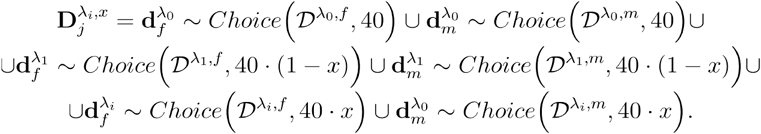

Furthermore, to be able to give an estimate of the variance in performance of our method per setting, we repeat this sampling process *n =* 10 times. This leaves us with a set of 10 datasets per log2 fold-change *l*_*i*_ and proportion of affected donors *x*:

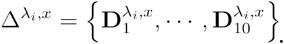

### Calculation of a normalized communication score per patient

To compare communication scores for a communication tetrad across patients, we derived a patient-specific communication score

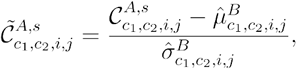

Where

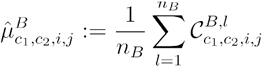

And

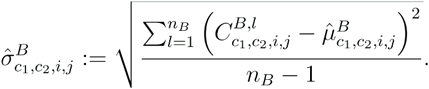

## Supplementary Figures

**Figure S1.**
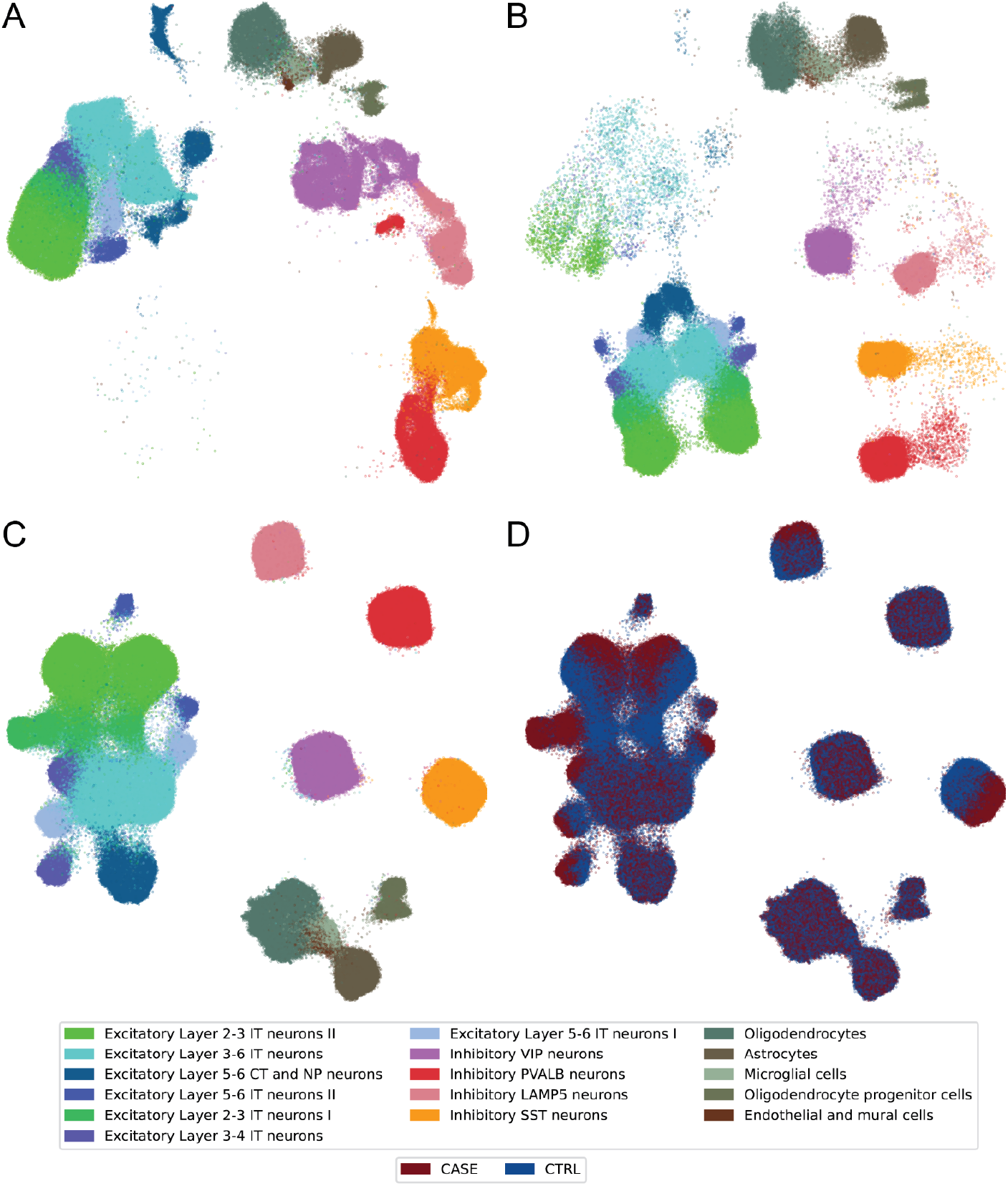
**A** Cell-type annotated Uniform Manifold Approximation Projection (UMAP) for all control cells from Bast et al. dataset. The projection was fitted in union with a set of simulated control cells. **B** Cell-type annotated UMAP for all cells from a cohort of 80 simulated control samples. The projections A and B reveal a distinct difference between simulated and original cells as the location in the latent space is highly different, while the rough structure between cell-types of the same subset (original vs. simulated) is preserved. **C** Cell-type annotated UMAP projection of all cells from a single simulated subset, including both control and case samples. **D** Same projection as in C, annotated according to the disease state association.

**Figure S2.**
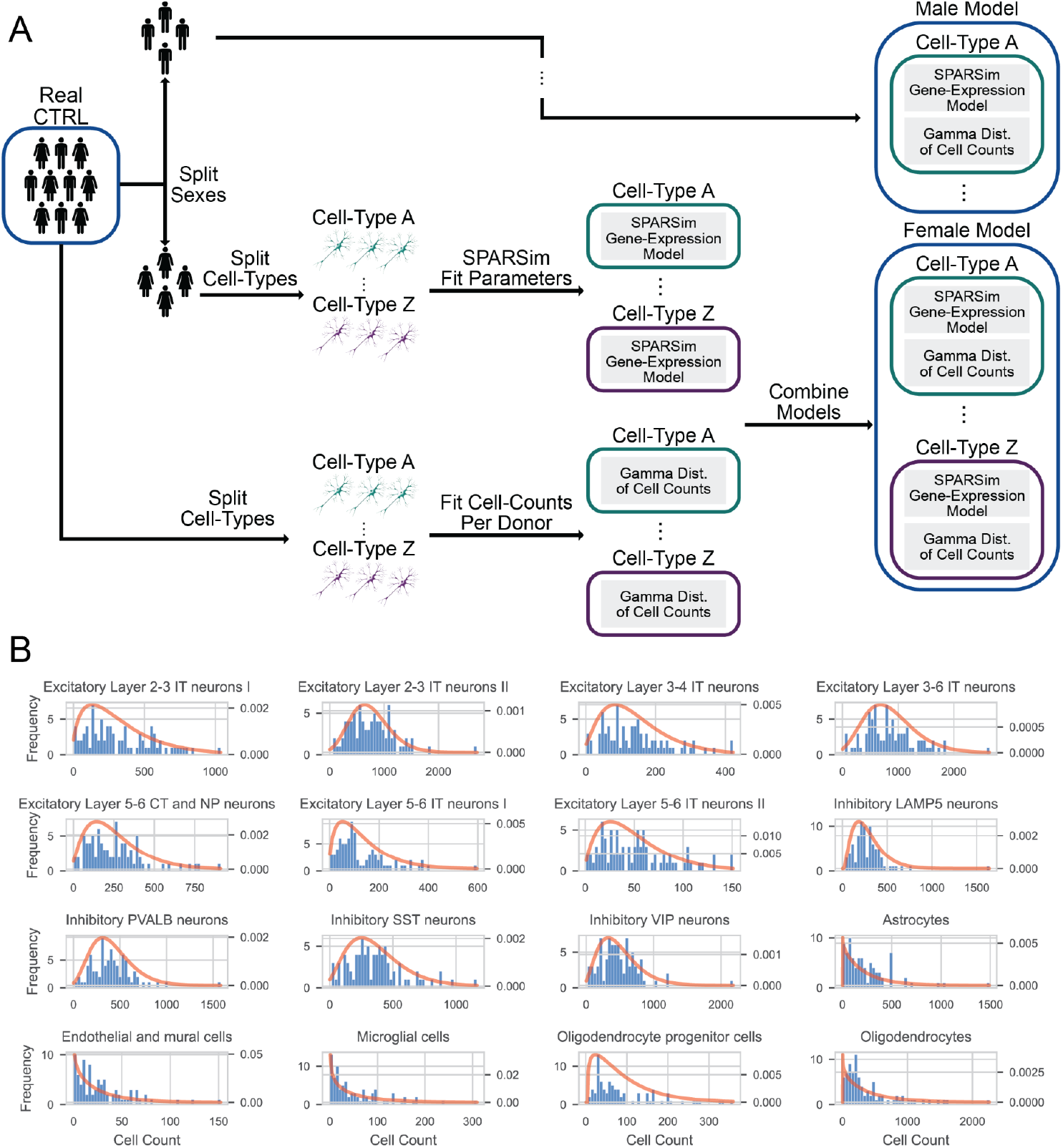
**A** Procedure for simulating the number of cells per donor using a gamma distribution (bottom) and transcription counts per cell-type using SPAR-Sim (Baruzzo, Patuzzi, and Di Camillo 2020). For simulating the transcription counts, we split the group by sexes and by cell-type to estimate gamma distribution parameters required for the simulation step. **B** Histograms of cell counts per cell type and donor (blue) and the fitted gamma distribution from which we sample the number of cells for a simulated subject (orange).

**Figure S3.**
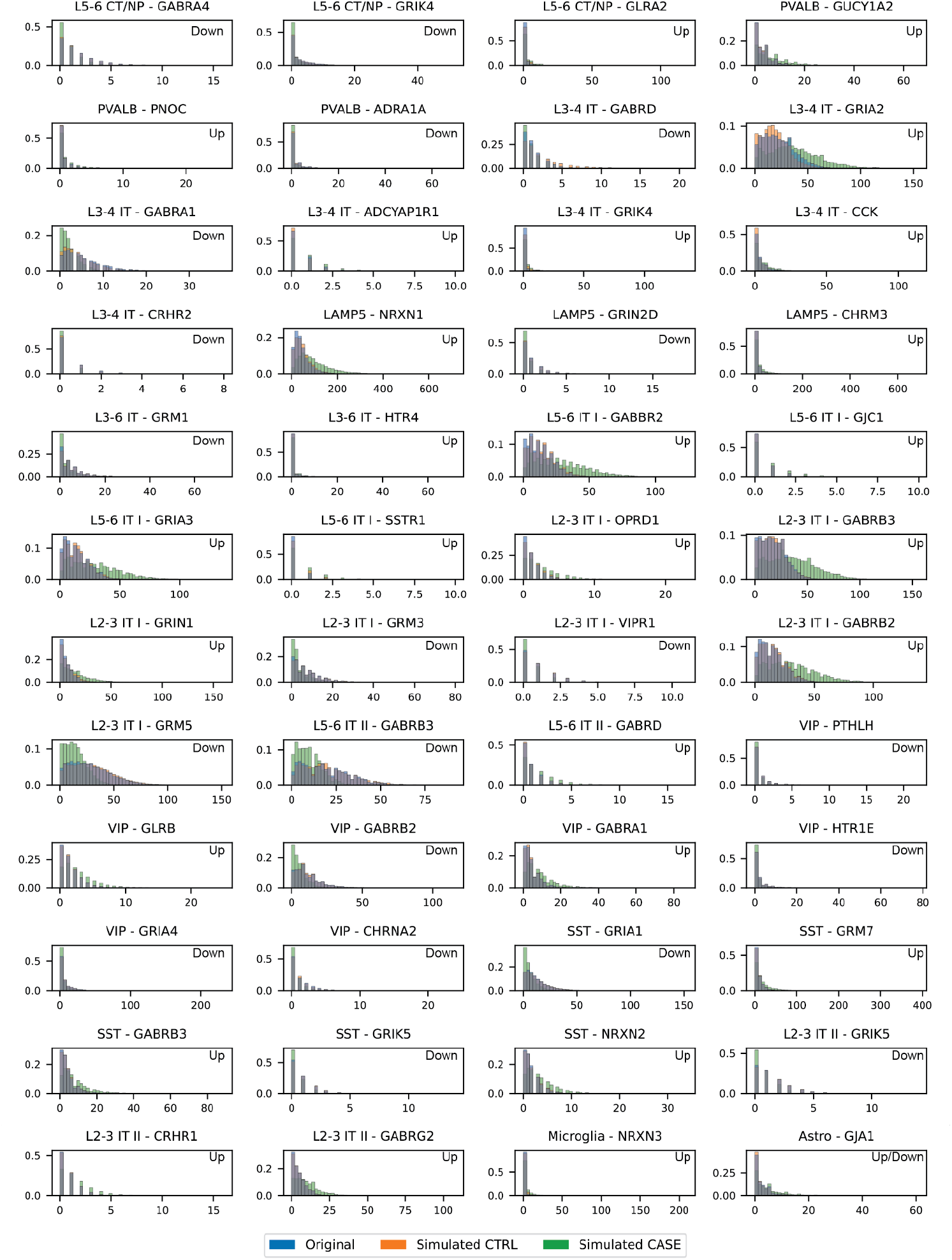
Histogram plot of raw gene expression counts for the original (blue), simulated CTRL (orange), and simulated CASE (green) dataset for cell types and genes that have been perturbed with a log_2_ fold-change of ±1.5 mean expression in the CASE group. As expected, gene expression of the original data and simulated unperturbed control group are matching, while changes in the gene expression of the simulated CASE group can be easily distinguished.

**Figure S4.**
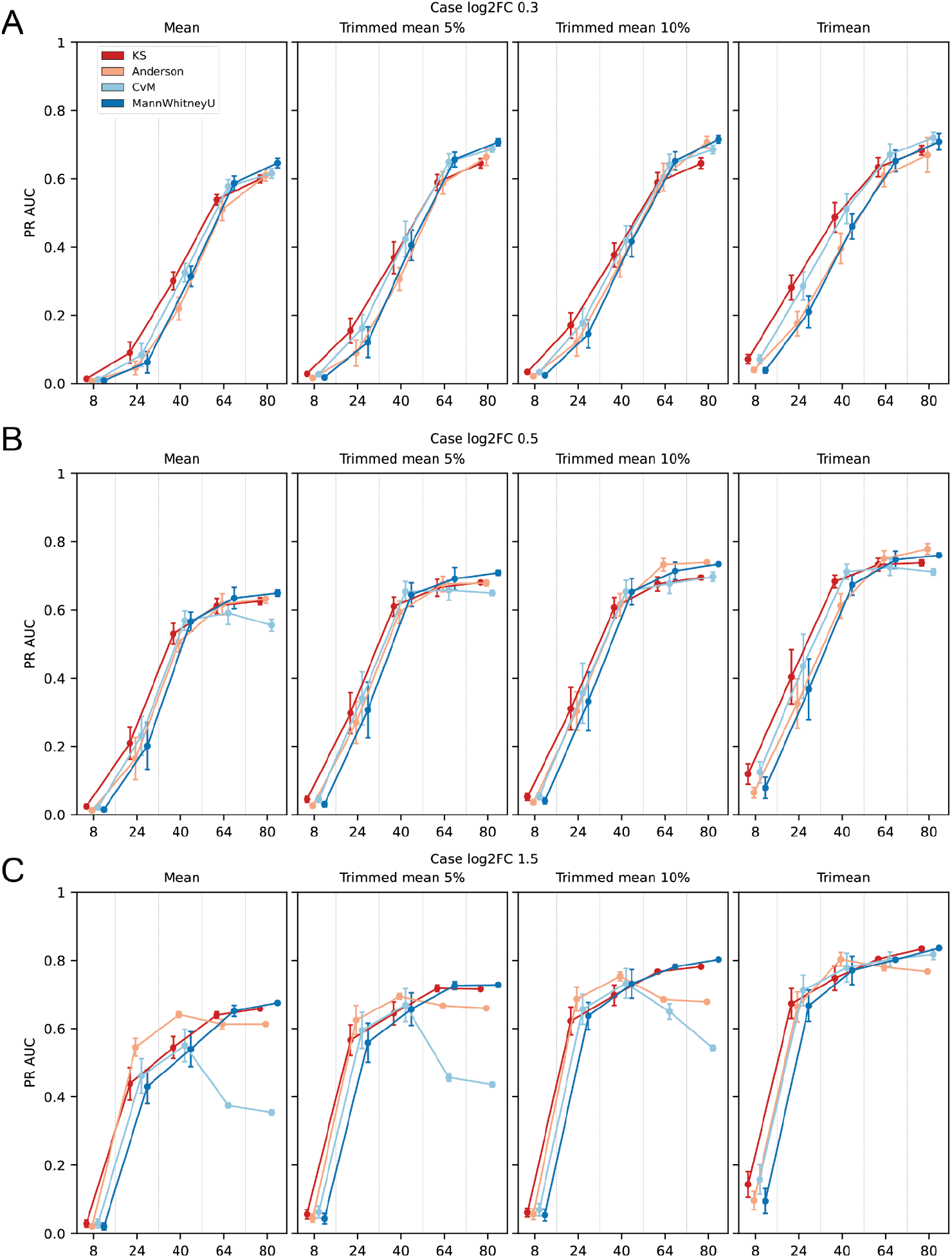
Comparison of the precision-recall area-under-the-curve (PR AUC) for the four different averaging modes, four different statistical tests, different proportions of affected donors (n out of 80), and different log_2_ fold-changes: **A** ±0.3, **B** ±0.5, **C** ±1.5.

## Tables

**Table T1:** MultiNeuronChat extended human database

T1_interaction_db_human.xsl

**Table T2:** Performed perturbations for in silico analysis

T2_perturbations.xlsx

**Table T3:** Runtime of MultiNeuronChat

T3_Runtime_Summary.xslx

**Table T4:** Results of in silico evaluation

T4_Simulation_Summary.xslx

**Table T5:** Results of ROSMAP analysis

T5_ROSMAP_summary.xlsx

## Data availability

The source data and preprocessed datasets are available through https://www.synapse.org/Synapse:syn3219045 (De Jager et al. 2018), and the European Genome-phenome Archive (Bast et al. 2025).

## Code availability

The MultiNeuronChat python package is available through GitHub https://github.com/Hjerling-Leffler-Lab/MultiNeuronChat and the code used to reproduce the analysis in this study is deposited at https://github.com/Hjerling-Leffler-Lab/MultiNeuronChat-Validation

## Acknowledgements

The authors are grateful for contributions to the MultiNeuronChat database: NeuronChatDB https://github.com/Wei-BioMath/NeuronChatAnalysis2022/tree/main/NeuronChatDB_table (Zhao et al. 2023) and the CellPhoneDB https://github.com/Teichlab/cellphonedb (Efremova et al. 2020). The results published here are in whole or in part based on data obtained from the AD Knowledge Portal (https://adknowledgeportal.org). Study data were generated from postmortem brain tissue provided by the Religious Orders Study and Rush Memory and Aging Project (ROSMAP) cohort at Rush Alzheimer’s Disease Center, Rush University Medical Center, Chicago. This work was funded by NIH grants U01AG061356 (De Jager/Bennett), RF1AG057473 (De Jager/Bennett), and U01AG046152 (De Jager/Bennett) as part of the AMP-AD consortium, as well as NIH grants R01AG066831 (Menon) and U01AG072572 (De Jager/St George-Hyslop). The simulation of in silico scRNAseq data is in part based on data obtained from (Bast et al. 2025).

The authors acknowledge support from the National Academic Infrastructure for Supercomputing in Sweden (NAISS)/ Uppsala Multidisciplinary Center for Advanced Computational Science for assistance with massively parallel sequencing and access to the PDC Center for High Performance Computing/ Uppsala Multidisciplinary Center for Advanced Computational Science (UPPMAX) computational infrastructure. Computational resources, data handling, data storage, and support were enabled by NAISS, hosted by UPPMAX and the PDC Center for High Performance Computing, partially funded by the Swedish Research Council through grant agreement no. 2022-06725.

## Author Contributions

Conceptualization (LB), Data Curation (GV), Formal Analysis (GV), Funding Acquisition (JHL), Investigation (GV, LB), Methodology (GV, LB), Project Administration (LB), Resources (JHL), Software (GV), Supervision (LB), Validation (all authors), Visualization (GV, LB), Writing - Original Draft Preparation (GV, LB), Writing - Review and Editing (all authors).

